# Working Memory Performance after Daily Caffeine Intake: Compromised Performance and Reduced Hippocampal Activity

**DOI:** 10.1101/2021.04.19.440520

**Authors:** Yu-Shiuan Lin, Janine Weibel, Hans-Peter Landolt, Francesco Santini, Helen Slawik, Stefan Borgwardt, Christian Cajochen, Carolin Reichert

**Affiliations:** Centre for Chronobiology, Psychiatric Hospital of the University of Basel, Switzerland; Transfaculty Research Platform Molecular and Cognitive Neurosciences, University of Basel, Switzerland; Neuropsychiatry and Brain Imaging, Psychiatric Hospital of the University of Basel, Switzerland; Institute of Pharmacology and Toxicology, University of Zurich, Switzerland; Sleep & Health Zurich, University Center of Competence, University of Zurich, Switzerland; Department of Radiology, Division of Radiological Physics, University Hospital Basel, Switzerland; Department of Biomedical Engineering, University of Basel, Switzerland; Clinical Sleep Laboratory, Psychiatric Hospital of the University of Basel, Switzerland

**Keywords:** Caffeine, withdrawal, working memory, hippocampus

## Abstract

Neuroprotective effects of caffeine have been frequently reported in the context of disease and cognitive dysfunction as well as in epidemiological studies in humans. However, evidence on caffeine effects on neural and memory functions during daily intake in a healthy cognitive state remains scarce. This randomized double-blind placebo-controlled crossover study investigated working memory functions by N-back tasks and functional magnetic resonance imaging (fMRI) after daily caffeine intake compared to a placebo baseline and to acute caffeine withdrawal in 20 young healthy volunteers. Each volunteer was given 3 times 150 mg caffeine for 10 days in the daily caffeine condition, 3 times 150 mg mannitol for 10 days in the placebo condition, and 9-day caffeine plus 1-day mannitol in the acute withdrawal condition. During the 10^th^ day, participants performed 4 N-back sessions (two loads each: 0- and 3-back) under controlled laboratory conditions. During the 4^th^ session of N-Back (i.e. at 5.5 h, 36.5 h and > 10 days after the last caffeine intake in the caffeine, withdrawal, and placebo condition, respectively) we assessed blood-oxygen-level-dependent (BOLD) activity. During the entire 10^th^ day, in 0-back tasks, we observed longer reaction times (RTs) in the withdrawal compared to the placebo (Cohen’s d = 0.7) and caffeine condition (Cohen’s d = 0.6), but no significant effects of conditions on error rates. In contrast, in 3-back tasks (controlled for 0-back), the RTs in the caffeine condition were longer compared to placebo (Cohen’s d = 0.6) and withdrawal (Cohen’s d = 0.5). Error rates were higher during both caffeine and withdrawal conditions compared to placebo (Cohen’s d of both contrasts = 0.4). Whole-brain analyses on fMRI data did not reveal significant condition-dependent differences in activities between task loads. Across task loads, however, we observed a reduced hippocampal activation (Cohen’s d = −1.3) during the caffeine condition compared to placebo, while no significant difference in brain activities between withdrawal and placebo conditions. Taken together, the worse working memory function and the hippocampal hypoactivation implicate a potential detrimental effect of daily caffeine intake on neurocognitive functions of healthy adults. Moreover, they echo the hippocampal volumetric reduction reported previously in the same volunteers. Lastly, acute withdrawal from daily caffeine intake impairs both low-order cognitive processes and working memory performance. Taking earlier studies on acute caffeine effects into account, our findings indicate that daily caffeine intake elicits a dynamic change in cerebral activities during the course of repeated consumption, with unknown consequences in the long run.

## Introduction

Caffeine is the most common and frequently consumed psychostimulant worldwide (Barone, 1984; Frary, Johnson, & Wang, 2005; Mitchell, Knight, Hockenberry, Teplansky, & Hartman, 2014). The benefit of acute caffeine intake on vigilance and mood has been repeatedly reported (Barry et al., 2005; Clark & Landolt, 2017; Urry & Landolt, 2015). Through the enhancement of simple and complex attention processes, caffeine can potentially benefit other higher-order cognitive functions (Clark & Landolt, 2017; Einother & Giesbrecht, 2013; Nehlig, 2010). Working memory function is crucial for high-order cognitive functions, and its performance also relies on basic low-order cognitive processes such as attention and motor controls (Diamond, 2013; Malenka RC, 2009). Thus, given its psychostimulation on attention and motor control, caffeine may enhance *apparent* performance in working memory tasks without eliciting *pure* influence on memory function (Nehlig, 2010). Acute caffeine intake was frequently found to improve working memory performance in shortening reaction time or when only assessing overall performances without detaching psychostimulation effects on the low-order cognitive processes (Haskell, Kennedy, Milne, Wesnes, & Scholey, 2008; Haskell, Kennedy, Wesnes, & Scholey, 2005; Morava, Fagan, & Prapavessis, 2019; Smith, Clark, & Gallagher, 1999). Studies, however, separating or statistically controlling for caffeine effects on low-order task performance (i.e. using high-against low-workload task to control for low-order processes) often reported no clear-cut net benefits on working or short-term memory function (García et al., 2017; Haller et al., 2017; Haller et al., 2013; Klaassen et al., 2013; Koppelstaetter et al., 2008; Schmitt, Hogervorst, Vuurman, Jolles, & Riedel, 2003).

Despite a devoid *pure* benefit to behavioral performance in working memory tasks, task-related brain responses towards acute caffeine seem to be more sensitive. Functional magnetic resonance imaging (fMRI) studies that examined blood-oxygen-level-dependent (BOLD) activities during working memory tasks and controlled for low-workload performance consistently reported an increase in regional activations after acute caffeine intake, yet without improving behavioral performances. Koppelstaetter et al. (2008) using 100 mg caffeine observed an acute increase in BOLD activity in medial frontal region without affecting N-back performance. Similarly, with 100 mg caffeine, Klaassen et al. (2013) also found an increased dorsolateral prefrontal activation in the encoding phase in a letter Sternberg task while a decreased thalamic activity in the maintenance phase, without significant effects on behavioral performance. Haller et al. (2013) using 200 mg caffeine reported an increase in task load-dependent BOLD activity in widespread cortical and subcortical regions in older healthy individuals, without significant behavioral differences in N-back performance. The same team later on compared the working memory performance after 200 mg caffeine between cognitive-stable and cognitive-declined elderly populations. Again, different post-caffeine activation in default mode network (DMN) between two groups was observed without significant behavioral effects (Haller et al., 2017). This evidence suggests that pharmacophysiological effects of caffeine could still modify memory-dependent brain functions without necessarily causing an apparent consequence in the behavioral performance of working memory tasks.

To date, a knowledge gap remains regarding the consequences of prolonging such cerebral effects by repeated daily exposure to caffeine, especially in a healthy cognitive state. Chronic caffeine intake has been frequently reported to rescue cognitive deficits induced by sleep loss, chronic stress, neurodegenerative diseases, and aging (Baur et al., 2020; Espinosa et al., 2013; Kaster et al., 2015; Laurent et al., 2014; Prediger, Batista, & Takahashi, 2005). These conditions, in fact, often embody a common characteristic of an upregulated expression of adenosine A2AR in the striatum (Cunha, 2016; Temido-Ferreira et al., 2020; Varani et al., 2010) and hippocampus (Canas, Duarte, Rodrigues, Köfalvi, & Cunha, 2009; Diógenes, Assaife-Lopes, Pinto-Duarte, Ribeiro, & Sebastião, 2007; Rebola et al., 2003; Viana da Silva et al., 2016). Adenosine modulates synaptic functions by controlling presynaptic neurotransmitter release and postsynaptic receptor activations through the counteraction between inhibitory A1 receptors (A1R) and excitatory A2A receptors [A2AR, (Ferre et al., 2005)]. Through chronic administration of caffeine, A1R, however, develops a tolerance to caffeine antagonism and attenuates its response, and the maintained effects of A2AR antagonism in turn prevails (Karcz-Kubicha et al., 2003; Quarta et al., 2004). Hence, the overexpressed adenosine A2AR may serve as a precondition for the profound counteraction of chronic caffeine against neuroexcitotoxicity with concomitant cognitive improvement, by reducing hippocampal presynaptic glutamate release (Martins et al., 2020), attenuating hippocampal synaptic long-term potentiation [LTP; (Costenla et al., 2011; Jerónimo-Santos et al., 2014; Lopes, Pliássova, & Cunha, 2019)], and preventing excitotoxicity induced by the hippocampal NMDA receptors antagonism (Dall’Igna et al., 2003; de Oliveira et al., 2005; Zhao et al., 2010). Yet, it remains unconcluded whether neural activities and memory functions in healthy individuals without specific circumstance-induced alterations can be negatively impacted by the reduced synaptic functions.

On the cognitive level, several observational studies have investigated cognitive effects of habitual caffeine intake but yielded mixed findings (discussed in (Lin et al., 2021)). Limited randomized controlled trials reported either no changes or clear tolerance to daily caffeine consumption in mood (Sigmon, Herning, Better, Cadet, & Griffiths, 2009), vigilance (Galduróz & Carlini, 1996; Judelson et al., 2005; Weibel et al., 2020), and memory-related performance (Galduróz & Carlini, 1996; Watson, Deary, & Kerr, 2002). On the cerebral level, while neurovascular effects were maintained over daily caffeine intake (Addicott, Peiffer, & Laurienti, 2012; Sigmon et al., 2009), our previous report revealed a concentration-dependent reduction of grey matter volumes in hippocampus after daily caffeine intake (Lin et al., 2021). These may mirror the aforementioned adaption in adenosinergic systems by long-term exposure to caffeine. Furthermore, daily caffeine intake can increase extracellular concentration of endogenous adenosine (Conlay, Conant, deBros, & Wurtman, 1997) and upregulate affinity of adenosine receptors for agonists (Jacobson, von Lubitz, Daly, & Fredholm, 1996; Varani et al., 2000; Varani et al., 2005; Varani et al., 1999). As a consequence, inhibitory adenosine signaling is strengthened when caffeine is ceased, resulting in cognitive and physiological withdrawal symptoms, such as increased drowsiness, difficulty in concentrating, flu-like symptoms, and elevated neurovascular dilation (Couturier, Laman, van Duijn, & van Duijn, 1997; Ferre et al., 2008; James, 1998; Juliano & Griffiths, 2004; Mathew & Wilson, 1985; Nikodijevic, Jacobson, & Daly, 1993; Sigmon et al., 2009; Weibel et al., 2020).

Hence, this double-blind randomized placebo-controlled crossover study aimed to investigate working memory function and its neural correlates after daily caffeine intake and during withdrawal. Based on the animal evidence on suppressed hippocampal excitation by maintained A2AR antagonism after chronic caffeine intake, we examined whether working memory performance and the underlying cerebral correlates in healthy humans could be negatively impacted during daily intake of caffeine. Furthermore, we expected that, through the upregulated A1R activation, acute caffeine withdrawal also led to a decreased task-related activities and impaired working memory performance. We measured working memory repeatedly across 13 hours after wake-up in a caffeine, a withdrawal, and a placebo condition using N-Back tasks. This procedure enabled to reliably map behavioral performance over the course of caffeine intake and metabolization, and the task allowed differentiating between “net” working memory function (i.e., controlled for basic psychomotor processed) and basic psychomotor process. While limiting treatment to the first 8 hours of each protocol, the fMRI scanning session took place after 13 hours to exclude acute short-term caffeine effects in BOLD activity patterns. Finally, we carefully controlled for caffeine-induced reductions in global perfusion in the functional analysis by statistically adjusting for whole-brain levels of cerebral blood flow.

## Methods

### Ethics

The ethical approval of the current study was issued by the Ethics Committee northwest/central Switzerland (EKNZ). The study execution followed the declaration of Helsinki, and all participants were fully informed with study details and consented in written form.

### Volunteers

Overall, twenty healthy male volunteers (age: 26.4 ± 4.0 years; Body mass index: 22.7 ± 1.4 kg/m^2^; and self-report habitual caffeine intake: 474.1 ± 107.5 mg/day) completed the study. Inclusion and exclusion criteria are addressed in detail in the supplement. One of the volunteer’s caffeine condition was excluded due to incompliance with the treatment.

### Study protocol

In a double-blind, randomized placebo-controlled study, each of the 20 volunteers completed three conditions: placebo, caffeine, and caffeine withdrawal (**Figure 1**). The order of the three conditions were randomized and semi-balanced (i.e. each order had three participants, while *Withdrawal – Placebo – Caffeine* and *Placebo – Caffeine – Withdrawal* had four participants). In each condition, volunteers underwent nine ambulatory days, followed by a 43-h laboratory stay, starting in the evening of the ninth day. Instead of washout periods, we implemented the 9-day ambulatory placebo to ensure a clean state off from remaining effects of caffeine and withdrawal (Juliano & Griffiths, 2004). It also ensure a standardized daily administration of caffeine with controlled dosage. On each day, volunteers received treatments at 45 min, 4 h, and 8 h after habitual rise time in the morning: In the placebo and the caffeine condition, the daily treatments were 3 times of 150 mg mannitol or 3 times of 150 mg caffeine plus additional mannitol, respectively, through 10 days; in the withdrawal condition, volunteers received caffeine capsules until the 9^th^ day, where they received caffeine for the first and placebo capsules for the second and third administrations until the end of tenth day. Actimetry-monitored rest-activity cycles were kept constant between conditions in timing and duration (8 h sleep, 16 h wakefulness) within each individual.

**Figure 1.**
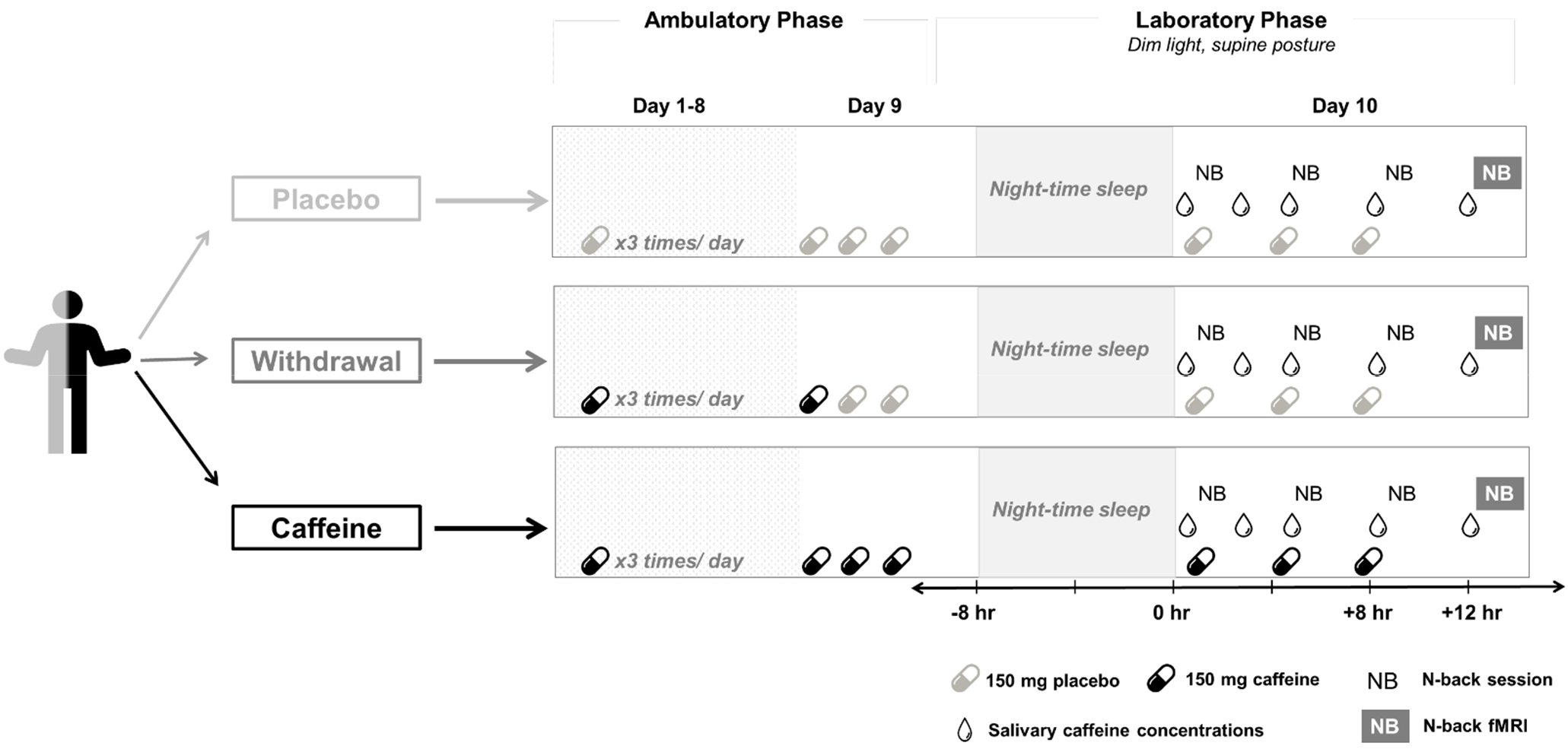
Study protocol.

Through the entire 10 days, volunteers abstained from caffeine-containing diets, including coffee, tea, energy drink, soda, and chocolate. The compliance was screened daily by collecting perspiratory caffeine levels in the evening [results see (Weibel et al., 2020)]. During the laboratory stay, volunteers stayed in dim light (<8 lux), constant half-supine position (~45°), with controlled dietary and lavatory time. Water consumption *ad libitum* did not differ between conditions [results see (Lin et al., 2021)]. Volunteers were not allowed to use mobile phones and had no social contacts except with the study personel.

Here, we focus on measurements (details in the following sections) aquired on the tenth day. All measurements were scheduled according to the individual’s habitual bedtime. The fMRI scan started at 13 h after habitual wake-up time (equivalent to 5 h after last treatment). Visual working memory tasks (N-back) were scheduled at 1 h, 5 h, 9 h, and 13 h (at scan) after awakening. We used salivary caffeine concentrations to ensure the successful administrations of the treatments.

### Measurements

#### 1. N-back

We employed N-back tasks to assess visual working memory on the laboratory day of each condition. Four sessions were scheduled during the course of the day and allowed us to study performance through and after caffeine intake. The timings of the four sessions relative to the treatments were: 15 min. after the first treatment, 1 h after the second, as well as 1 h and 5 h after the last treatment. Every session consisted of 9 trials of 3-back and 5 trials of 0-back, each trial consisted of 30 stimuli with 1.5 s interval in a quasi-random order. Each stimulus was presented for 1s. The stimuli were letters, which were presented consecutively on a screen. Participants answered with key “1” when the current letter matched the one three stimuli earlier (3-back, “high load”) or when it was a “K” (0-back, “low load”), otherwise, pressing “2”. For habituation, participants performed one practice session in the evening of the ninth day.

Performances were indexed by (a) error rate, calculated from the ratio of “incorrect rate” (missed + false alarms) to “correct response” (hits + correct rejections), and (b) reaction time (RT) in correct answers. We employed generalized linear mixed models to analyze the condition and time effects by using R packages afex (Henrik Singmann, 2019) and lme4 (Bates, Mächler, Bolker, & Walker, 2015). We defined “working memory function” by the porportion of the performance in 3-back against 0-back. To examine the net working memory performance in 3-back, we regressed out the variance of 0-back. Furthermore, we included the order of conditions as a covariate in order to control for learning effects and their potentially confound with the condition effect. Distinct from our previous report (Lin et al., 2021), the current analyses sepcifically focused on N-back performances in 3 conditions with 4 sessions each. During scanning, task presentation was identical as during the other sessions except that participants reacted with an fMRIcompatible response box. Particpants were asked to react with index (for “1”) and middle finger (for “2”) irrespecitve of using a keyboard or a reponse box.

#### 2. Functional magnetic resonance imaging (fMRI)

##### Acquisition

We used a two-dimensional multislice gradient-echo echo-planar imaging sequence (GRE-EPI, 3 x 3 x 3 mm^3^, TR = 2500 ms, TE = 30 ms, FA=82°, number of slice = 39) to measure task-related blood-oxygen-level-dependent (BOLD) activities, and a three-dimensional magnetization-prepared rapid gradient-echo (MP-RAGE) sequence (1×1×1mm3, TR=2000 ms, TI= 1000 ms, TE=3.37ms, FA=8°) to acquire T1-weighted structural images on a 3T Siemens scanner (MAGNETOM Prisma; Siemens Healthineer, Erlangen, Germany).

##### Image preprocessing

We employed the preprocessing pipeline for EPI and T1-weighted images from CONN toolbox (http://www.nitrc.org/projects/conn) on Matlab. The preprocessing of both structual and functional images started with spatial realignment, unwarping, and corrections for slice-timing and motions (motion threshold set at 0.9mm). Structural and functional images were normalized to standard brains from Montreal Neurological Institute (i.e. MNI-space) independently, followed by the co-registration and the segmentation of grey matter (GM), white matter (WM), and cerebrospinal fluid (CSF). The images were then resliced into 2 mm and 1 mm isotropic voxels for functional and structural data, respectively. Finally, the functional images were smoothed with Gaussian kernel of 8 mm full width half maximum.

##### Classical functional analysis

The statistical analysis was performed on SPM12. In the first level analysis, we used 3- and 0-back as regressors of interests and six estimated parameters of motion as regressors of no interests. Signal drifts were adjusted by a high-pass filter of 128s and serial correlations using an AR(1) model. We contrasted the activation by task-loads, i.e. 3-back > 0-back, 3-back < 0-back, and load-independent responses. In the group-level analysis, we used a flexible factorial model to estimate condition effects as the fixed effect, subject effects as the random effect, and age as a covariate, in each task-load contrast. In order to control for the caffeine- and withdrawal-induced changes in perfusion, we used individual whole-brain cerebral blood flow, measured by arterial spin labelling (sequence detail see the *Acquision* section in methods of (Lin et al., 2021)), as a covariate of ANCOVA for global normalization. The statistical significant thresholds were set at a voxel-level uncorrected p < 0.001, and at a cluster-level threshold set at P_FWE_ < 0.05.

##### Functional connectivity analysis

As the functional connectivity analysis was an exploratory step based on the observations in the cluster analysis, we employed a ROI-to-ROI approach and focused on the hippocampus and middle frontal gyri. The ROIs were defined and segmented based on the FSL Harvard-Oxford atlas (including 132 ROIs).

We performed a functional connectivity analysis using the CONN toolbox. Before the classical first level analysis, a denoising step using the anatomical component-based noise correction procedure (aCompCor) was implemented. We used linear regressions to remove the potential confounding effects in the BOLD signal, including the noise signals derived from WM and CSF, the estimated motions, the identified outlier scans (i.e. scrubbing), and the constant and first-order linear session effects. Further, we applied a temporal band-pass filter of 0.008 – 0.09 Hz to reduce the low-frequency drift derived from physiological sources or head-motions.

The first level analysis was performed with a weighted general linear model with bivariate correlations. We used Hemodynamic Response Function (HRF) to weigh down the beginning for a delay of the BOLD response. For the group analysis, we used two task-loads and three conditions as regressors of interest and age as regressor of no interest. The statistical significant thresholds were set at a voxel-level uncorrected p < 0.001, and at a cluster-level threshold set at P_FWE_ < 0.05.

#### 3. Salivary caffeine levels

We collected salivary samples in 2-h intervals from 15 min before the first treatment intake until the end of the laboratory day in each condition. As a validation for successful treatments, we focused on the five samples before caffeine administration, 1 h after each administration, and 15 min before the scan. The salivary caffeine concentrations were quantified by a High Performance Liquid Chromatography (HPLC) coupled to tandem mass spectrometry at the Laboratory Medicine, University Hospital Basel. The chromatographic separation was done by an analytical ion exchange phase column for methyl malonic acid. The detection threshold of the method was 20 ng/ml. We analyzed the sample x condition effets on caffeine levels by generalized linear mixed models with R packages afex (Henrik Singmann, 2019) and lme4 (Bates et al., 2015).

## Results

### Caffeine levels

The salivary caffeine data confirmed a successful experimental manipulation among placebo, caffeine, and withdrawal conditions. A significant condition x sample effect (F_2,91_ = 17.7, p < .001) indicated that, from baseline to time of scan, a significant increase in concentrations in the caffeine condition and a significant decreased in concentration in the withdrawal condition, while no significant change in caffeine concentrations in placebo condition.

**Table 1.** Salivary caffeine concentration (μg/ml) per condition. Mean and standard deviation of the samples collected before the first treatment as well as before the fMRI scan session are presented per condition. “Baseline” indicates the first sample in the morning of the laboratory (10^th^) day before administration of scheduled caffeine or placebo. The asterisks indicate the samples showing significant increase from baseline to the time of scan compared to placebo.

|  | Placebo | Caffeine | Withdrawal |
| --- | --- | --- | --- |
| Baseline | 0.03 ± 0.03 | 1.18 ± 1.24 | 0.46 ± 0.82 |
| Time of scan | 0.03 ± 0.04 | 3.67 ± 2.08 *** | 0.12 ± 0.18 ** |

\*\* $: p < .01$ ; \*\*\* $: p < .001$

### Cognitive performance

#### Basic attentional and psychomotor performance (o-back)

On the 0-back error rates, we did neither observe a significant effect of condition (F_2,198_ = 1.9, p = .152, **Figure 2**) nor a significant condition x session interaction (F_6,198_ = 1.4, p = .216). Independent of condition, we found a significant session effect (F_3,198_ = 5.1, p = .002), for which a post-hoc analysis indicated a worse performance in Session 2 and 3 compared to both Session 1 (p_all_ < .040, Cohen’s d = 0.46 and 0.49) and scan session (p_all_ < .007, Cohen’s d = 0.57 and 0.59). [Median (quartile) of error rates in 0-back: placebo 0.02 (0.01 - 0.03); caffeine: 0.02 (0.01 – 0.04); withdrawal 0.03 (0.02 – 0.04)].

**Figure 2.**
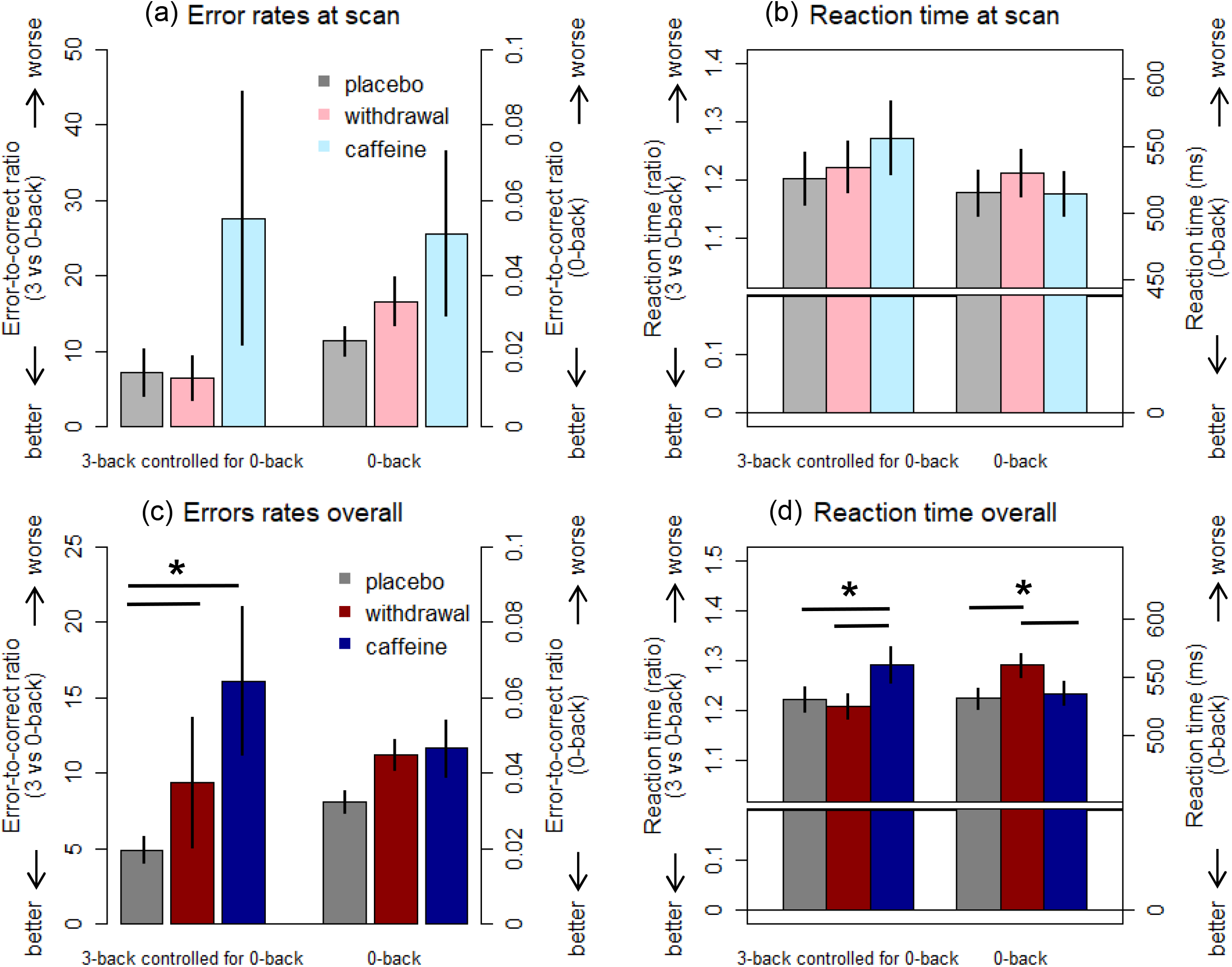
Main effect and time course of N-back performance per condition. **(a)** and **(b)** display the main effects of conditions <u>at scan session</u> with means and standard errors of the error rates (left panel) and the reaction time (RT, right panel). The left side of each plot presents the performance in 3-proportional to 0-back, while the right side of each presents the performance in 0-back tasks. In the identical fashion, (c) and (d) display the main effects of conditions <u>over all four sessions</u>. ***Note:*** the proportions of 3- to 0- back here are <u>only to visualize</u> the adjustment of the variance. In statistics, we used raw data of 3-back as the outcome variables and regressed out the variance of 0-back in a linear mixed model.

On the RTs in 0-back, we observed a significant effect of condition (F_2,198_ = 9.6, p < .001). A post-hoc analysis yielded a longer RT in the withdrawal condition compared to both the placebo (p < .001, Cohen’s d = 0.68) and caffeine conditions (p = .003, Cohen’s d = 0.58). No significant difference was found between caffeine and placebo in RTs (p = .815). Moreover, we found a significant session effect (F_3,198_ = 8.3, p < .001), for which the post-hoc analysis indicated exceptionally shorter RTs in the scan session compared to all other sessions (p_all_ < .011, Cohen’s d = −0.61 – −0.92). No significant interaction between condition and session was found (F_6,198_ = 0.5, p = .790). [Mean ± SD of RTs (ms) in 0-back: placebo 531.7 ± 71.4; caffeine: 536.1 ± 85.6; withdrawal 560.3 ± 83.5].

Performance in 0-back at the scanning session is separately presented in **Figure 2**. As summarized above, the interactions of session x condition were, however, not significant on error rates nor on RTs. Therefore, we have no indication for a specific condition-dependent modulation of performances at the scan session.

#### Working memory function (3-back controlled for o-back)

On the *net* error rates of 3-back, we observed a significant main effect (F_2,197_ = 5.0, p = .008). A post-hoc analysis indicated a higher *net* error rate in the caffeine (p < .031, Cohen’s d= 0.40) and withdrawal (p = .013, Cohen’s d= 0.45) condition compared to placebo. In **Figure 2**, we illustrate this effect by presenting the ratio of 3-back to 0-back per condition. The analyses did not indicate a significant difference between caffeine and withdrawal conditions (p = .960). No significant main effect of session (F_3,195_ = 1.4, p = .244) nor condition x session interaction (F_6,195_ = 0.3, p = .936) was found. [Median (quartile) in 3-back: placebo 0.05 (0.03 - 0.09); caffeine: 0.07 (0.03 - 0.14); withdrawal 0.07 (0.03 - 0.14)].

Similarly, we also found a significant main effect of condition on *net* 3-back RTs (F_2,196_ = 6.6, p = .002), in which a significant longer *net* RT was observed in the caffeine condition compared to both placebo (p = .003, Cohen’s d= 0.57) and withdrawal (p = .023, Cohen’s d= 0.47). No significant difference was found in *net* 3-back RTs between withdrawal and placebo. In addition, we observed a significant session effect (F_3,195_ = 3.4, p = .019), in which the prominent difference was a significantly longer RT in the first compared to the last sessions (p = .022, Cohen’s d= 0.60). We did not find a significant condition x session interaction (F_6,196_ = 0.2, p = .960) in the RTs. [Mean ± SD of RTs (ms) in 3-back: placebo 647.5 ± 128.2; caffeine: 688.9 ± 179.0; withdrawal 671.2 ± 123.6].

Again, in order to properly interpret the following fMRI results, we separately present the 3-back performance (error rate and RTs) at the scan session in **Figure 2**. We emphasize that the analyses presented above, however, did not reveal any significant interactions of session x condition. Therefore, we did not have no clear-cut indications of condition-dependent differences in the performance at scan sessions specifically.

### Functional analysis

Independent of conditions, the whole brain analysis indicated a significant difference in the BOLD activity between 3- and 0-back (**Figure 3**). In particular, we observed increased activities in the bilateral attention and motor regions in 3-back compared to 0-back, including premotor and supplementary motor areas, parietal lobes, and superior temporal gyrus. Moreover, we found reduced activities in 3-back compared to 0-back in DMN regions and parts of the limbic system, including left posterior cingulate cortex, left medial prefrontal cortex, bilateral angular gyri, bilateral medial temporal gyri, and bilateral hippocampi. These load-dependent activities did, however, not show a significant difference among the three conditions.

**Figure 3.**
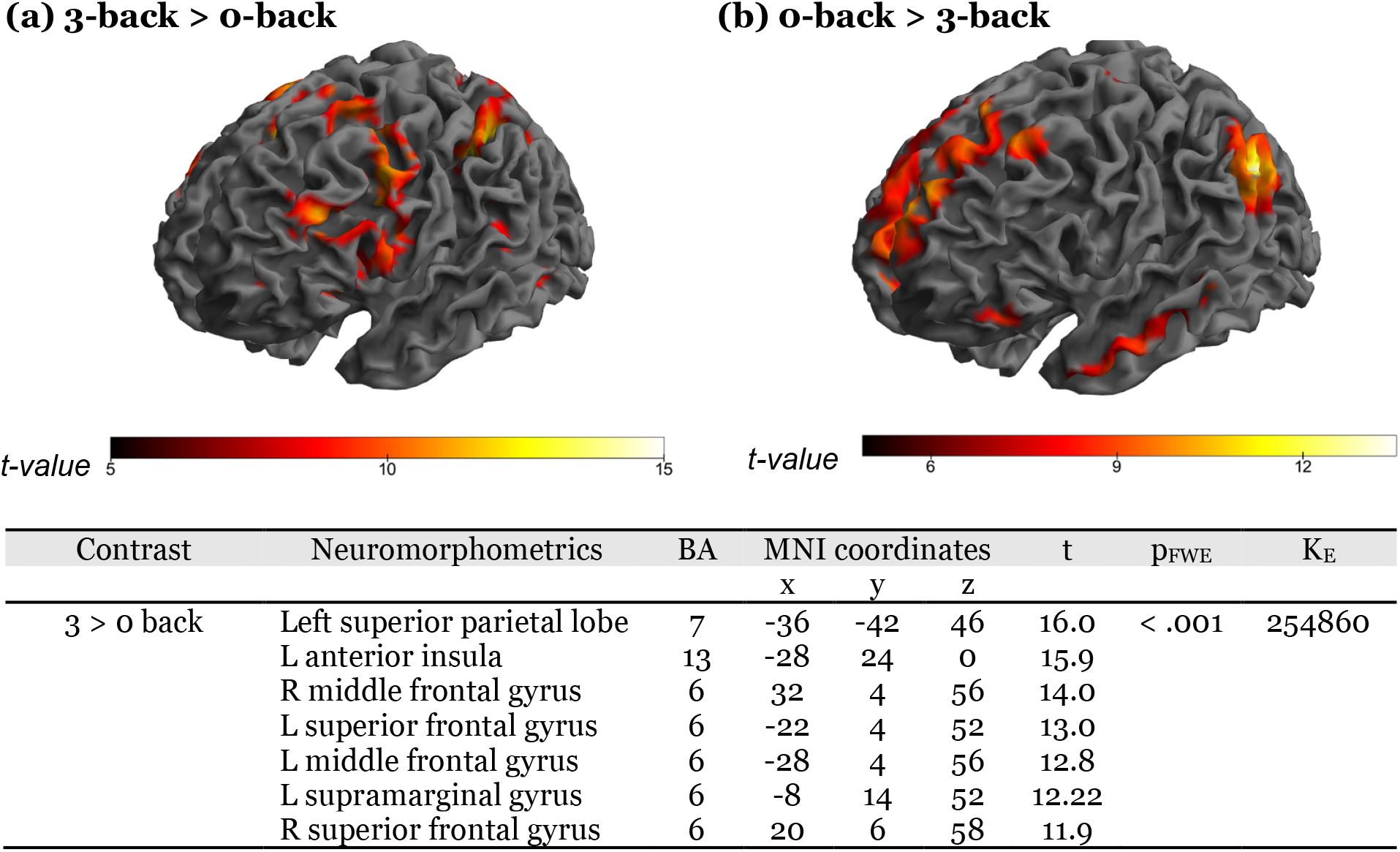
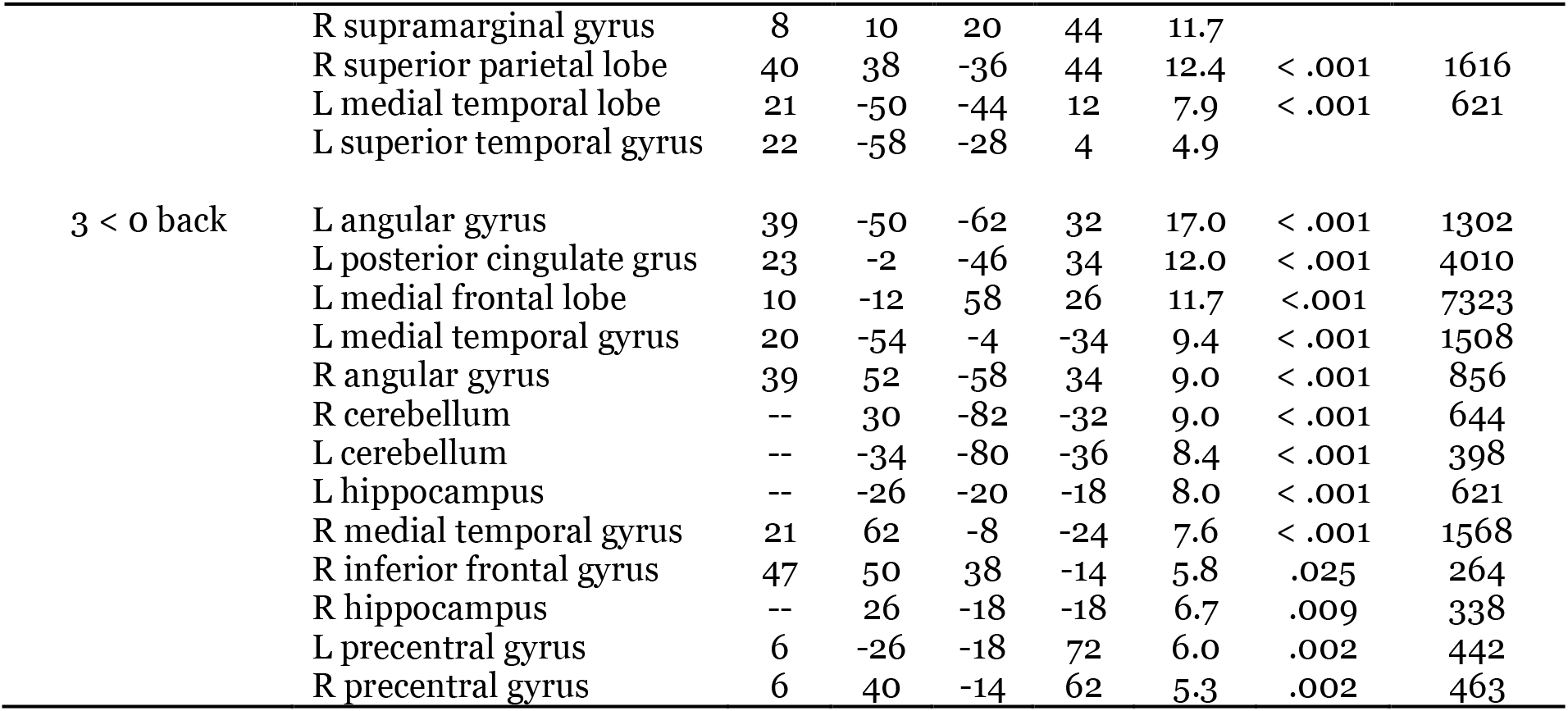
Task load-related blood-oxygen-level-dependent (BOLD) activity, independent of conditions. **(a)** brain regions showing increased activities during 3-back compared to 0-back. **(b)** brain regions showing decreased activities during 3-back compared to 0-back. ***Table:*** The table presents the SPM results, including peak level t-values, cluster level p_FWE_ values, and cluster sizes (K_E_). We also denote MNI coordinates of each region and their corresponding atlas labels based on neuromorphometrics and the Brodmann’s area (BA) from whole-brain analysis. Multiple brain regions covered in a large cluster (such as left SPL and all in 3 > 0 back) are displayed with shared cluster level statistics.

Irrespective of task loads, we observed an overall lower BOLD activity in caffeine condition in the right hippocampal region (p = .035, Cohen’s d= −1.3, detailed statistics see **Figure 4**) and an at-trend (p_FWE_ = .057, Cohen’s d= 1.2) higher activity in left middle frontal gyrus, compared to placebo. Moreover, compared to withdrawal, we also found a higher activity in the right middle frontal gyrus in the caffeine condition (p < .001, Cohen’s d= 1.1). No significant differences in the regional activation was observed between withdrawal and placebo.

**Figure 4.**
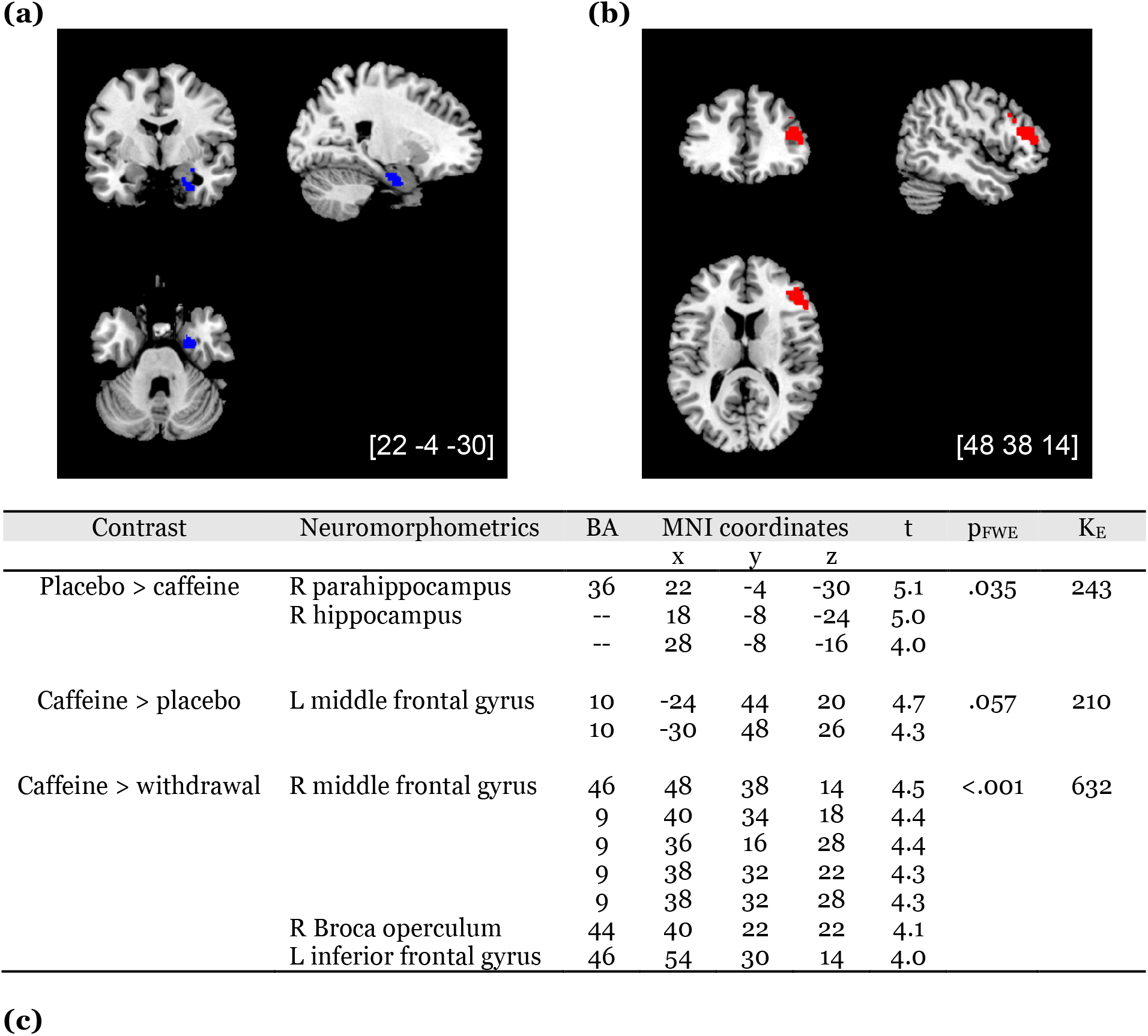
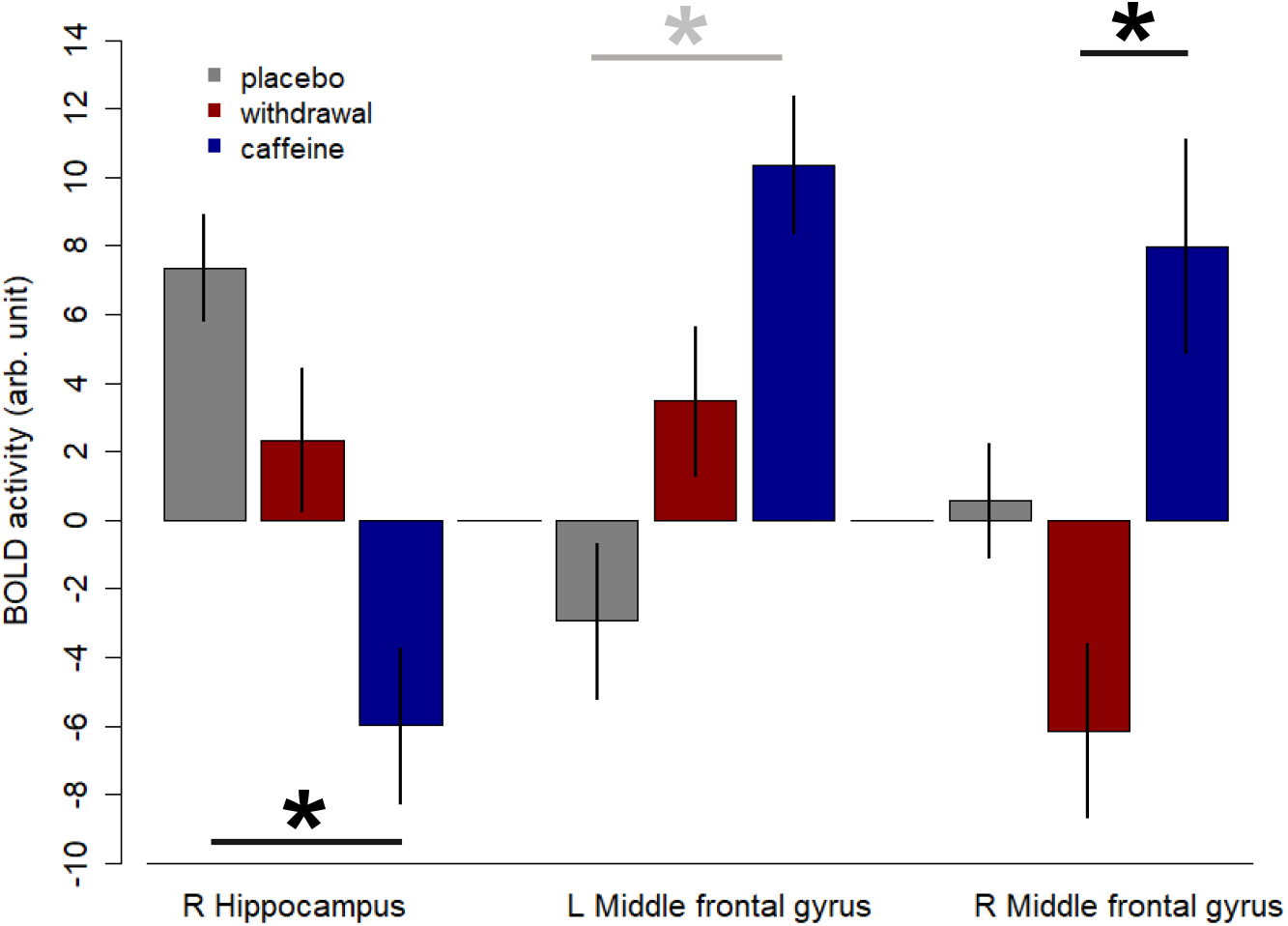
Location and direction of task-related BOLD activtiy differences between conditions. **(a)** the blue color indicates the location of reduced right hippocampal BOLD activity (p_FWE_ < .05) during daily caffeine intake compared to placebo. **(b)** the red color indicates the location of increased BOLD activity (p_FWE_ < .05) in the right middle frontal gyrus during daily caffeine intake compared to withdrawal. ***Table:*** The table presents the SPM results, including peak level t-values, cluster level p_FWE_ values, and cluster sizes (K_E_). We also denote MNI coordinates of each region and their corresponding atlas labels based on Neuromorphometrics and the Brodmann’s area (BA) from whole-brain analysis. Multiple brain regions covered in a large cluster (such as right parahippocampus *et al.* in placebo > caffeine) are displayed with shared cluster level statistics **(c)** Eigenvariates of the significant clusters in hippocampus and left and right middle frontal gyri. The black asterisks and black lines indicate the statistically significant contrast, while the grey asterisk and the grey line indicate the contrast only exhibited at trend differences.

Based on the reduced hippocampal activity and the *at-trend* increased activation in middle frontal gyrus, we hypothesized a caffeine-strengthened anticorrelation between these two regions. We exploratorily examined the functional connectivity between bilteral hippocampus and middle frontal gyri. However, no significant differences were found among conditions, irrespective of task-loads.

## Discussion

Here we investigated working memory performance and the underlying cerebral correlates after 10 days of regular caffeine intake and during caffeine withdrawal, compared to a 10-day placebo-controlled condition. The absence of caffeine effects on reaction time in 0-back, yet a slower reaction time in the withdrawal condition, indicate a tolerance that may have developed towards the classical caffeine effects on enhancing low-order cognitive processes. On the contrary, working memory functions were compromised after daily caffeine consumption and after withdrawal. The typical patterns of functional brain activity in attention, motor, and default mode networks in working memory, however, did not differ after caffeine intake comapred to placebo. Independent of working memory challenges, we found an universally reduced BOLD activity in the right hippocampus after daily intake compared to placebo, which echos our previously reported caffeine-induced hippocampal grey matter reduction (Lin et al., 2021). Adding to the literature, our study indicates a detrimental impact of daily caffeine intake on brain functionality in healthy volunteers, including a compromised working memory function and a reduced hippocampal activity. Compared with the neuroprotective evidence of chronic caffeine intake in disease or cognitive dysfunction models, our findings suggest divergent effects of caffeine in a healthy and a deficient cognitive state.

The performance of N-back involves hierarchical cognitive processes comprising multiple components, which heavily depend on different task loads. 0-back, a simple recognition-response task, involves mostly attention, motor control, and inhibtion processes without challenging memory capacity; 2-or more back requires components recruited in 0-back plus an increasing memory capacity and continuous information updating. The current study revealed a worsened low-order process in the withdrawal condition but no differences during daily caffeine intake, compared to placebo. This finding corresponds to our previous report in a psychomotor vigilance task, where we observed a potential tolerance during daily caffeine intake but a deteriorated attention during withdrawal (Weibel et al., 2020). In acute administration protocols (i.e. when caffeine is administered after a phase of abstinence in a “clean” state), caffeine elicits the classical enhancement of mood, attention, and motor control by facilitating adenosine-modulated dopaminergic and excitatory signaling (Ferre, 2008, 2010; Kaasinen, Aalto, Någren, & Rinne, 2004; Volkow et al., 2015). However, adenosinergic properties can be adpated over daily intake, including altered gene expression of receptors (Svenningsson, Nomikos, & Fredholm, 1999), upregulated affinity of receptors for agonists (Jacobson et al., 1996; Varani et al., 2000; Varani et al., 2005; Varani et al., 1999), elevated concentration of endogenous adenosine (Conlay et al., 1997), and reduced binding or expression to antagonists (Kaplan, Greenblatt, Kent, & Cotreau-Bibbo, 1993; Karcz-Kubicha et al., 2003). Such adaptions attenuate effects of acute caffeine challenge and incurs a tolerance (Karcz-Kubicha et al., 2003; Newland & Brown, 1997; Svenningsson et al., 1999). Moreover, in acute caffeine withdrawal, the strengthened adenosine activation could exert stronger inhibition such as diminishing dopaminergic signals (Ferre, 2010, 2016), resulting in poor attention and motor disinhibition (Nieoullon, 2002; Nieoullon & Coquerel, 2003; Ott & Nieder, 2019).

The remaining puzzle is that working memory functions were worse after caffeine intake and did not recover during the first 36 hour of withdrawal. During the daily intake of caffeine when A1R develops tolerance, behavioral effects of caffeine are primarily accounted for by the antagonism to the non-tolerant A2AR prevails (Ferre, 2010; Ferre et al., 2008). In hippocampus, A2AR controls the presynaptic glutamate release, while A1R modulates the inhibition of NMDA receptors (Martins et al., 2020; Rebola, Canas, Oliveira, & Cunha, 2005). Thus, daily caffeine intake may lead to a synaptic hypoactivity in hippocampus by constantly inhibiting glutamate release without commensurate availability of posynaptic NMDA receptors. Furthermore, through the maintained antagonism to hippocampal A2AR, daily caffeine attenuates hippocampal longterm potentiation (LTP) in healthy freely-behaving rodents (Blaise, Park, Bellas, Gitchell, & Phan, 2018). LTP a marker of persistent synaptic strength which is considered to be one of the cellular mechanisms of memory and learning (Bliss & Collingridge, 1993; Cooke & Bliss, 2006). Supported by our data indicating reduced hippocampal activities in caffeine compared to placebo, it is possible that, the synaptic hypoacitivites progressively influence the hippocampal strength and function over daily intake of caffeine, leading to alterations in the hippocampal structures, a universally compromised activities, and suboptimal working memory functions.

To date, substantial evidence using disease models or derived in a deficient cognitive state has shown an established neuroprotective effect of caffeine against cognitive impairments and synaptic dysfunctions induced by age, stress, and sleep loss (Baur et al., 2020; Espinosa et al., 2013; Kaster et al., 2015; Laurent et al., 2014; Prediger et al., 2005). Compared to the evidence in a compromised physiological state, our data suggest a potential divergence of daily caffeine effects in a healthy state on working memory performance and cerebral activity. Nevertheless, the observed effects in our study may additionally be attributed to the specific dose we administered, or even to the specific habitual intake levels of our participants. First of all, given the inverse-U dose effects of caffeine, high-dose caffeine could inversely jeopardize cognitive functions, trigger excitotoxicity, and potentiate apoptosis (Fredholm, Bättig, Holmén, Nehlig, & Zvartau, 1999; Xie, Huang, Li, Wang, & Huang, 2021). Although the current study used a dose roughly equivalent to three times of double espresso (Heckman, Weil, & Gonzalez de Mejia, 2010) to simulate a daily repeated patterns of caffeine intake, a couple of studies using equivalent to lower caffeine doses with chronic administration in rodents [dose conversion: 10 mg/kg in rodents were estimated to be equivalent to 250 mg/70 kg in humans (Fredholm, Battig, Holmen, Nehlig, & Zvartau, 1999)] have reported a suppressed neurogenesis in healthy adult hippocampus and concomitant impairment in long-term memory (Han et al., 2007; Wentz & Magavi, 2009). Furthermore, daily intake was found to decelerate the elimination rate of caffeine compared to acute administration (Lau, Ma, & Falk, 1995), and daily administration of a higher dose further disproportionates the accummulation of its primary metabolite, paraxanthine (Denaro, Brown, Wilson, Jacob, & Benowitz, 1990). Thus, over daily intake, concentrations of the active metabolites may progresisvely increase, exceed the safety limit, and increase the risk to induce detrimental effects.

In addition, participants in our study had a relatively high amount of habitual intake, which may have exerted a precondition for the responses observed in the study. More clinical studies should further investigate dose responses of hippocampal functionality and of working memory functions to daily caffeine intake.

In addition, we also observed a lower BOLD activity in the medial frontal gyrus (encompassed in dorsolateral prefrontal cortex, DLPFC) in withdrawal compared to caffeine condition. The DLPFC is critical for cognitive functions involving goal-oriented attention processes (Bahmani et al., 2019; Clark, Squire, Merrikhi, & Noudoost, 2015). The deviant in DLPFC activity in both task loads between withdrawal and caffeine conditions may implicate a universally less and more engagement of attention-related regions, respectively. In line with our earlier report on a worse vigilance across the day after caffeine withdrawal compared to caffeine, this observed difference in DLPFC activity mirrors the cerebral susceptibility of attentional processes to the abrupt cessation of daily caffeine intake (Weibel et al., 2020).

Irrespective of conditions, our functional MRI data corroborated the increased activity in the attention and motor regions (including prefronatal, parietal, and temporal cortices), as well as reduced activity in DMN regions (including posterior cingulate cortex, medial prefrontal cortex, angular gyrus, and medial temporal lobe), in 3-back task relative to 0-back. Comparing with the previous evidence which showed increased brain activities without changing behavioral performance after acute caffeine intake (Haller et al., 2017; Haller et al., 2013; Klaassen et al., 2013; Koppelstaetter et al., 2008), the absence of condition effects on load-dependent cerebral responses in our data might point out a certain tolerance effect. Moreover, in comparison to earlier studies, our scan session was located at a time of day exerting strong circadian wake-promotion, which may have interacted with the effects of condition on cerebral responses (Byrne, Hughes, Rossell, Johnson, & Murray, 2017; Reichert et al., 2017). Behaviorally, we observed an improved low-load performance at the time of the scans compared to other sessions, which could mask the effects of condition on the load-dependent behavioral and cerebral responses.

The current study bears a few strengths compared to the literature. The most significant impact is to first demonstrate the impacts of daily caffeine intake and caffeine withdrawal on working memory function on both behavioral and cerebral levels. In addition, taking advantage from the existing measurement for cerebral blood flow, we could statistically control for the caffeine- and withdrawal-induced responses of brain perfusions (Addicott et al., 2012; Addicott et al., 2009; Sigmon et al., 2009). Moreover, the study adopted a strictly controlled clinical trial design to standardized the daily exposure to caffeine, i.e. precise dose, durations of treatment, monitored abstinence. We used a corssover design to minimize the bias from sequential effects. We applied 10 days of placebo intake with monitored abstinence to confirm the washout of remaining caffeine levels and avert manifestations of withdrawal symptoms in the measurement of baseline. In additon, to standardize the effect of duration of wakefulness on the efficacy of caffeine, we scheduled the scans, treatments, tasks, and sleep according to the individual habitual bedtime. During the laboratory stay, the physical activities and diets of the volunteers were standardized, and the amount of water consumption was recorded. These features allow precise attributions of observed effects to the treatments.

On the other hand, a few features of our study could limit the data interpretation and should be carefully considered. First of all, caffeine can induce neurovascular uncoupling by reducing baseline CBF and increasing baseline cerebral metabolic rate of oxygen consumption (CMRO2) (Perthen, Lansing, Liau, Liu, & Buxton, 2008). Although we mitigated the deviation by statistically adjusting for the variance of resting cerebral blood flow, to resolve the issue more precisely, future studies are recommended to adopt a sequence supporting spiral readouts to obtain simultaneous CBF and BOLD activties (Perthen et al., 2008). In addition, despite a crossover design, the analysis might be still restricted with relatively small sample size, due to two missing scans (one in placebo and one in caffeine). Furthermore, in order to reduce variances within small sample size, our data suffered from a lack of diversity in sex differences, therefore the generalizability in female population is limited. Moreover, as the current study used unified dosage in a population of moderate to high habitual consumers, the effects of different daily doses on the healthy population with lower habitual intake should be further investigated. Lastly, the impacts observed in a crossover design entails that the effects observed in daily caffeine intake can also be restored in maximal 10 days (i.e. the duration of placebo administration prior to the placebo scan).

In conclusion, our data suggested a potential detrimental effect of daily caffeine intake on working memory function. Moreover, the reduced hippocampal BOLD activity after daily caffeine intake adds up to the previous evidence on daily caffeine-induced grey matter decrease in hippocampus. Our findings indicate an impact of daily caffeine intake in healthy adults, divergent from the evident acute effects and from the neurocognitive enhancement in deficient cognitive state. However, the crossover design also implicated the restorability of such alterations. Hence, more longitudinal studies employing a longer follow-up period with controlled treatments are required to examine whether the observed changes will further adapt or be maintained. Moreover, in light of the contrary outcomes currently observed in a healthy cognitive state as compard with earlier evidence on deficient cognitive functions, we recommend more studies to shed lights on the molecular mechanisms underlying daily caffeine effects in young and healthy habitual caffeine consumers.

## Acknowledgement

We express our sincere appreciation to our interns Andrea Schumacher and Laura Tincknell, M.Sc. student Sven Leach and Joshua Kistler, as well as all the study helpers for their assistance in the experiment and data processing. We also thank Dr. Corrado Garbazza and Dr. Martin Meyer for the health check during screening process. We are grateful for the assistance and resources provided by Professor Katharina Rentsch and Dr. Sophia Rehm at the Laboratory Medicine, University Hospital Basel. We especially appreciate all our participants for their volunteering and cooperation. Conflicts of interest: No conf licts of interest to declare.

